# In for a penny, in for a pound: Examining motivated memory through the lens of retrieved context models

**DOI:** 10.1101/464388

**Authors:** Deborah Talmi, Deimante Kavaliauskaite, Nathaniel D. Daw

## Abstract

When people encounter items that they believe will help them gain reward, they later remember them better than others. A recent model of emotional memory, the emotional Context Maintenance and Retrieval model (eCMR), predicts that these effects would be stronger when stimuli that predict high and low reward can compete with each other both during encoding and retrieval. We tested this prediction in two experiments. Participants were promised £1 for remembering some pictures, but only a few pence for remembering others. Their recall of the content of the pictures they saw was tested after one minute and in Experiment 2, also after 24 hours. Memory at immediate test showed effects of list composition. Recall of stimuli that predicted high reward was greater than of stimuli that predicted lower reward, but only when high- and low-reward items were studied and recalled together, not when they were studied and recalled separately. More high-reward items in mixed lists were forgotten over a 24-hour retention interval compared to items studied in other conditions, but reward did not modulate the forgetting rate, a null effect which should be replicated in a larger sample. These results confirm eCMR’s predictions, although further research is required to compare that model against alternatives.

## Introduction

It is paramount to remember information that can help us attain our goals. Because memory resources are limited, it is adaptive for the brain to prioritise experiences that can be leveraged to maximise subjective utility over less-important experiences (Gershman & Daw, 2017). This intuition was expressed by Sherlock Holmes, Arthur Conan Doyle’s timeless protagonist, when, justifying his ignorance of astronomy, he remarked that “It is of the highest importance… not to have useless facts elbowing out the useful ones”. Holmes’ intuition agrees with rational considerations, which suggests that people should allocate their limited resources to increase recall of the most valuable information (Anderson & Schooler, 1991). Yet how the opportunity to maximise utility influences the cognitive mechanisms that underlie successful recall has not been fully worked out.

There is substantial evidence from classical and instrumental conditioning tasks that the immense capacity of encoding (Bjork & Bjork, 1992) are biased towards subjectively-valuable information. For example, stimuli that predict reward attract preferential attention (e.g. Austin & Duka, 2010; reviewed in Le Pelley et al., 2016 and Mather, Clewett, Sakaki, & Harley, 2016). Since attention is one of the key drivers of subsequent memory, it is, perhaps, not surprising that in an experimental setting where participants anticipate that remembering some items earns more reward, operationalised through money or ‘points’, such items are rehearsed more frequently (Stefanidi et al., 2018) and remembered better (Ariel & Castel, 2014; Hennessee et al., 2019; Lee et al., 2015; Stefanidi et al., 2018; Wittmann et al., 2005). This bias is partially automatic, evident in that reward can attract attention obligatorily, even against current goals (Bucker & Theeuwes, 2017), and high-reward items that participants are immediately instructed to forget are nevertheless recognised more accurately (Hennessee et al., 2019). Nevertheless, there is also an element of top-down control on this bias, evident in that it is possible for participants to attenuate it by allocating resources to non-rewarding items (Hennessee et al., 2019). The bias reward exerts on the allocation of encoding resources clearly agrees with the broad rational analysis of memory, mentioned above. Yet while the computational and neural aspects of motivated memory have been discussed extensively (Lisman et al., 2011; Mason et al., 2017; Mather et al., 2015; Rouhani et al., 2018; Shohamy & Adcock, 2010), less is known about the cognitive mechanisms (Barsalou, 2017) that process signals associated with reward anticipation to support memory formation, maintenance and retrieval. Our main objective in this paper, therefore, is to investigate how reward anticipation influences memory-relevant cognitive mechanisms.

In this endeavour, it may be helpful to look to the emotional memory literature for clues. Theories of emotion define it as a response to signals that are appraised as goal-relevant (Lazarus, 1991; Sander et al., 2005). From this perspective, both a picture scene that depicts violent crime and a handshake that clinches a business deal reward can be described as emotionally significant. Like reward, emotional stimuli also attract preferential attention (Pourtois et al., 2013), modulate consolidation (Cahill & McGaugh, 1998) and enhance memory (for review, see Ack-Baraly et al., 2016). Although the definition of emotion is controversial (Barrett, 2006; Izard, 2010), as is the boundary between emotion and motivation (Chiew & Braver, 2011), it is useful to consider whether they influence memory through the same cognitive mechanisms.

The emotional Context Maintenance and Retrieval model (eCMR) is a variant of retrieved-context models, which was developed to account for the influence of emotion on free recall (Talmi et al., 2019). eCMR models emotion reductively through a single parameter, which was introduced to capture the preferential processing resources typically allocated to emotional stimuli (Pourtois et al., 2013). In addition to their effect on processing resources during encoding, emotional and neutral stimuli are also thought to differ in the pattern of semantic associations among them. However, given that in reward experiments there is no reason to be concerned with particular structures of semantic associations, it is the influence on attention that will be key here. eCMR distinguishes between two aspects of the encoding context – the temporal and the source context (Polyn et al., 2009). The ‘temporal context’ is a core concept for retrieved-context theory (Howard & Kahana, 2002). The term does not refer to the passage of time per se (Hintzman, 2016), but rather to the sequential nature of encoding, whereby the encoded representation of each event reflects its place in the sequence of events. Thus, the temporal context of each item is a recency-weighted representation of previous items. By contrast, the ‘source context’ codes the attributes that one item shares with others regardless of their place in the sequence. In an experiment, the source context refers to aspects of encoding outside its identity. Stimuli such as emotional picture scenes trigger unique cognitive, affective and physiological changes (Bradley et al., 1992), and therefore, in emotional memory experiments, the source context of emotional items is emotional while that of neutral items is neutral. In eCMR (compared to previous retrieved-context models) the extra attention that participants allocate to emotional items binds them more tightly to their source context. When memory is probed with a free recall test, the temporal context of the test allows participants to narrow their memory search to that of the most recent study set, thereby minimising intrusions (Lohnas et al., 2015). When items from the previous set compete with each other for recall, the one whose encoding context is most similar to that of the test context is the most likely to be recalled (Polyn et al., 2009), and its recall biases the test context, thereby influencing the recall likelihood of subsequent items. The tighter binding of emotional items to the emotional source context will enhance their competitive advantage during the test when they compete against neutral items (Talmi et al., 2019), while making it difficult to link them to a new context later on (Madan et al., 2012). This is how eCMR explains the robust findings of superior recall of distressing scenes and taboo words in typical experimental settings, where emotional and neutral stimuli are presented and recalled together, in ‘mixed’ lists. eCMR also successfully predicts a less intuitive finding – that the recall advantage of emotional items is less pronounced and sometimes disappears altogether when emotional and neutral items are presented and recalled separately, in ‘pure’ lists (Barnacle et al., 2017; Hadley & MacKay, 2006; MacKay et al., 2004; Talmi et al., 2007). The joint pattern of recall of emotional and neutral stimuli in mixed and pure lists is called the emotional list composition effect. eCMR simulates this effect as a consequence of the interplay between encoding – where emotional items always capture attention, regardless of the composition of the list, and are therefore more tightly bound to their source context - and retrieval effects, where the increased association strength between emotional items and the source context helps them win the retrieval competition only against neutral items, but not against equally strongly-bound emotional items. The reader is referred to previous publications for the formal definition of these ideas (Howard & Kahana, 2002; Lohnas et al., 2015; Polyn et al., 2009; Talmi et al., 2019).

Although we developed eCMR to simulate the effects of emotion, the model makes the same prediction for experiments that manipulate reward, for two reasons. First, as aforementioned, both emotional and high-reward items attract processing resources preferentially. Second, the unique aspects of processing either of these item types – such as appraisal of goal relevance and goal congruence, physiological and phenomenological arousal - can be construed as rendering their ‘source context’ as unique and distinguished from the neutral source context of non-emotional or low-reward stimuli. For these reasons, eCMR makes the same prediction for a list-composition effect for reward as it does for emotion. This prediction is somewhat counter-intuitive, because it seems adaptive for memory to be best for items that predict “a pound” over those that predict “a penny”.

eCMR unequivocally predicts that the promise for higher gain would only result in a recall advantage in the mixed list condition, but other theoretical positions are more equivocal. While it is manifestly adaptive to only retain valuable information, rational analysis may suggest that it could be even more adaptive to retain all information unless there is a cost to doing so – even if this contradicts the haughty retorts from 221B Baker Street. Rationale considerations may induce participants either to allocate preferential resources only to high-reward items (resulting in a main effect of reward, and a null effect of list type), or to all but the low-reward items in mixed lists (resulting in a list-composition effect). Theoretical positions that account for the list-strength effect – essentially, the effect of list composition effects on recall of items that are strengthened through spaced repetition - predict null effects of list composition when items are strengthened by capturing preferential encoding resources (Malmberg & Shiffrin, 2005). Finally, theories that focus on the influence of reward at the maintenance stage (Lisman et al., 2011; Shohamy & Adcock, 2010) are often silent about immediate effects, although they predict a main effect of reward on delayed recall.

So far, eCMR has only been tested against results of experiments with negatively-valenced emotional materials. We are not aware of any previous studies of the list-composition effect with positively-valenced stimuli, nor with items that predict reward explicitly and thus may also be perceived as positively-valenced. List-composition effects have been reported for other types of materials that attract special elaboration during encoding (McDaniel & Bugg, 2008), and for stimuli that are repeated multiple times within a study set (Ratcliff et al., 1990), but not for stimuli that are studied for longer durations, for mass-repeated items, and for deeply-encoded items (Malmberg & Shiffrin, 2005). Here we test the prediction of eCMR, that reward will give rise to the list composition effect, for the first time.

We tested this hypothesis by examining recall of picture stimuli whose later recall explicitly predicts high or low reward, and which were encoded in pure and mixed lists. In two experiments, our participants viewed lists of neutral pictures that were presented framed or unframed. They knew that they could gain larger monetary reward by recalling framed items on later test. The critical manipulation was the arrangement of framed and unframed items in the list, such that some lists included both item types (mixed) and some only one (pure). The frame provides a clear signal of which individual stimulus should be prioritised (Mather et al., 2016; Mather & Sutherland, 2011). The main aim of Experiment 2 was to provide a conceptual replication of Experiment 1 by repeating the procedure in a new sample, with small changes in methodology. It also included a delayed test of the same effect to explore whether the delayed effect of list composition would be modulated by reward.

## Results

### Experiment 1

#### Simulation results

Because our predictions are based directly on simulations using eCMR, we begin by using eCMR to simulate Experiment 1. The simulations reported here use the same formal structure described previously (Simulation 2, Talmi et al., 2019). Notably, a post-encoding distractor is already included in that simulation in the manner typical for retrieved-context models, simulating additional studied items that cause temporal context drift but cannot be recalled. The main difference between previously-reported and new simulations is the semantic associations of stimuli in mixed lists. In typical emotional memory experiments the emotion manipulation is implemented by introducing a single ‘emotional’ category and a single ‘neutral’ category in each of the mixed lists, thereby creating a particular pattern of semantic associations (see, for example, Figure 1, middle panel). By contrast, in the experiments reported here and most motivated memory experiments the reward manipulation does not impact upon listwise semantic association strengths. In keeping with emotional memory experiments, in Experiment 1 mixed lists included two semantic categories (we use the values 0.08 and 0.02 here for within- and between-category associations, respectively), while in Experiment 2 they included only one semantic category.”

**Figure 1.**
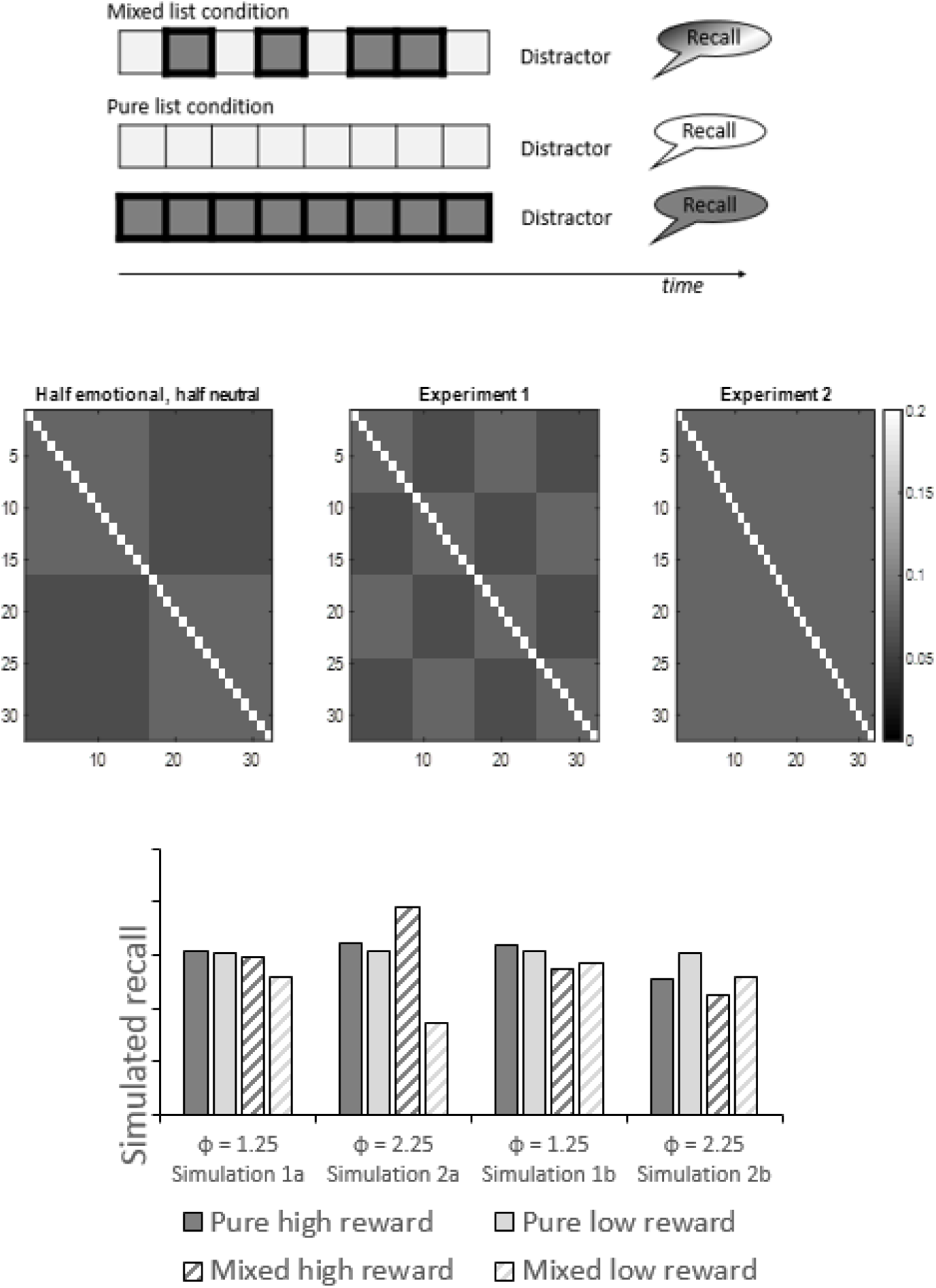
Model simulations of the list-composition task. Top. Schematic of the list composition task. The list composition task comprises of 2 types of lists: pure lists with items of only one type, and mixed lists of items of two types. Traditionally, one type of items is ‘stronger’ in some way. Here the two item types were framed pictures that predicted £1 if recalled and unframed pictures that predicted £0.10 if recalled. These are depicted as squares with thicker and thinner frames in the figure. At the end of list presentation participants recall all of the items they studied in any order. Here, a brief distractor task intervened between encoding the recall. Middle. Graphical representation of the hypothetical strength of semantic associations between stimuli in one mixed list in the current Experiments 1 and 2, and, for comparison, in typical emotional memory experiments. Each row/column represents a single item in the list. Darker shades of grey represent stronger semantic associations. The leftmost matrix models Talmi et al., 2007. Half of the stimuli are negative, arousing picture scenes (here, the first half rows/columns) and the other half, neutral domestic scenes. The middle matrix models Experiment 1. This experiment also included two semantic categories in each list, crossed with the reward manipulation. The first quarter of the stimuli are the framed clothing pictures, followed by framed stationery pictures, unframed clothing, and unframed stationery. The rightmost matrix models Experiment 2, where there was only a single semantic category in each list. Although half the items were framed here, as well, this manipulation did not change their simulated semantic strength.”. Bottom. Results of two simulations of Experiment 1 using eCMR.

The parameter used previously to model the increased attention to distressing emotional scenes was used here to model the increased attention to framed pictures that promised higher reward, where a value of *φ_emot_* = 1 represents a null effect of reward on attention. Simulation 1 used *φ_emot_* = 1.25 (the value of this parameter in Simulation 2 of Talmi et al., 2019), and Simulation 2 *φ_emot_* = 2.25 to expose the effect that differential motivational investments might have on recall. All parameter values are reported in Tables 1 and 2. eCMR simulates the influence of item type on encoding resources by allowing *φ_emot_* to modulate the strength of the context-to-item association (it modulates 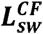 ). Simulations 1b and 2b examine an alternative, where *φ_emot_* modulate the rate with which the temporal context is updated (to modulate 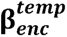 ).

**TABLE 1.**
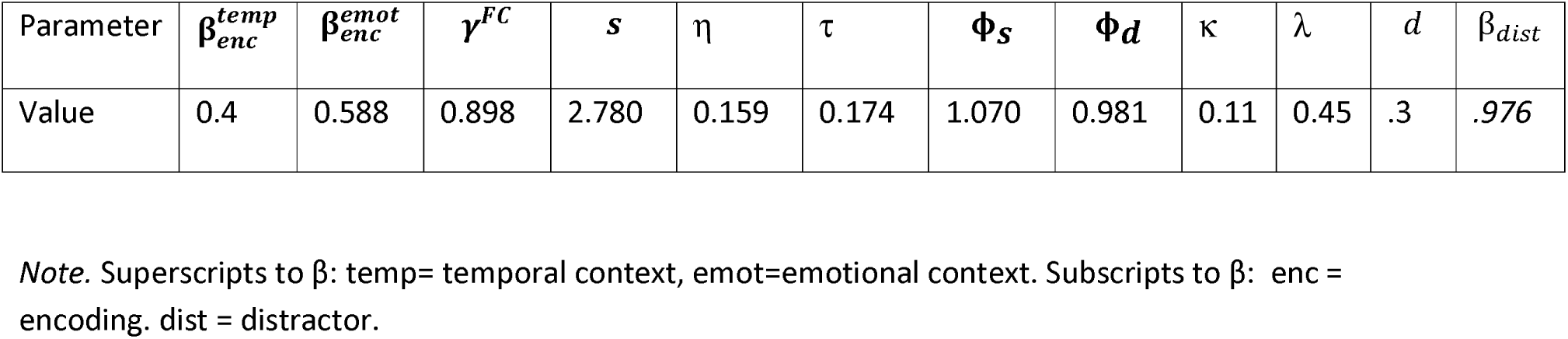
Fixed parameter values in simulations.

**TABLE 2.** Variable parameter values in Simulations 1-5.

| Parameter | *Talmi et al. 2019* | 1 | 2 | 3 | 4 | 5 | 6 |
| --- | --- | --- | --- | --- | --- | --- | --- |
| $\phi_{emot}$ | 1.25 | 1.25 | 2.25 | 2.25 | 2.25 | 2.25 | 2.25 |
| $\beta_{rec}^{temp}$ | .51 | .51 | .51 | .51 | .05 | .05 | .05 |
| $\beta_{rec}^{emot}$ | .588 | .588 | .588 | .588 | .588 | .05 | .05 |
| $L_{tw}^{CF}$ | .3 | .3 | .3 | .3 | .3 | .3 | .1 |
| $L_{wt}^{FC}$ | .898 | .898 | .898 | .898 | .898 | .898 | .1 |

*Note.* Superscripts to L: CF = context to item; FC: item to context; Subscripts to L: tw = temporal context to item; wt = item to temporal context. emot = emotional, rec = recall, dist = distractor.

The bottom panel of Figure 1 depicts the results of the simulations. Simulations 1a and 2a show even when the list comprises of two semantic categories, as in Experiment 1, the model still predicts that the reward manipulation will elicit a list composition effect. Increasing the potency of the reward manipulation (by increasing *φ_emot_* in 2a compared to 1a) only influenced the mixed list condition, where it increased the recall of high-value items and decreased the recall of low-value items. Comparing figures 1 and 2 show that the lower value of *φ_emot_* in simulation 1a provided a better qualitative fit to the average recall data than the value used in simulation 2a. By contrast, simulations 1b and 2b did not capture the emotional list composition effect; the increased drift between items led to *decreased* recall of the in mixed lists in comparison to pure lists. When the effect of emotion was made stronger (by increasing *φ_emot_* in 2b compared to 1b), the increase in contextual drift between items did not decrease the recall of emotional items as much as that of neutral items.

**Figure 2.**
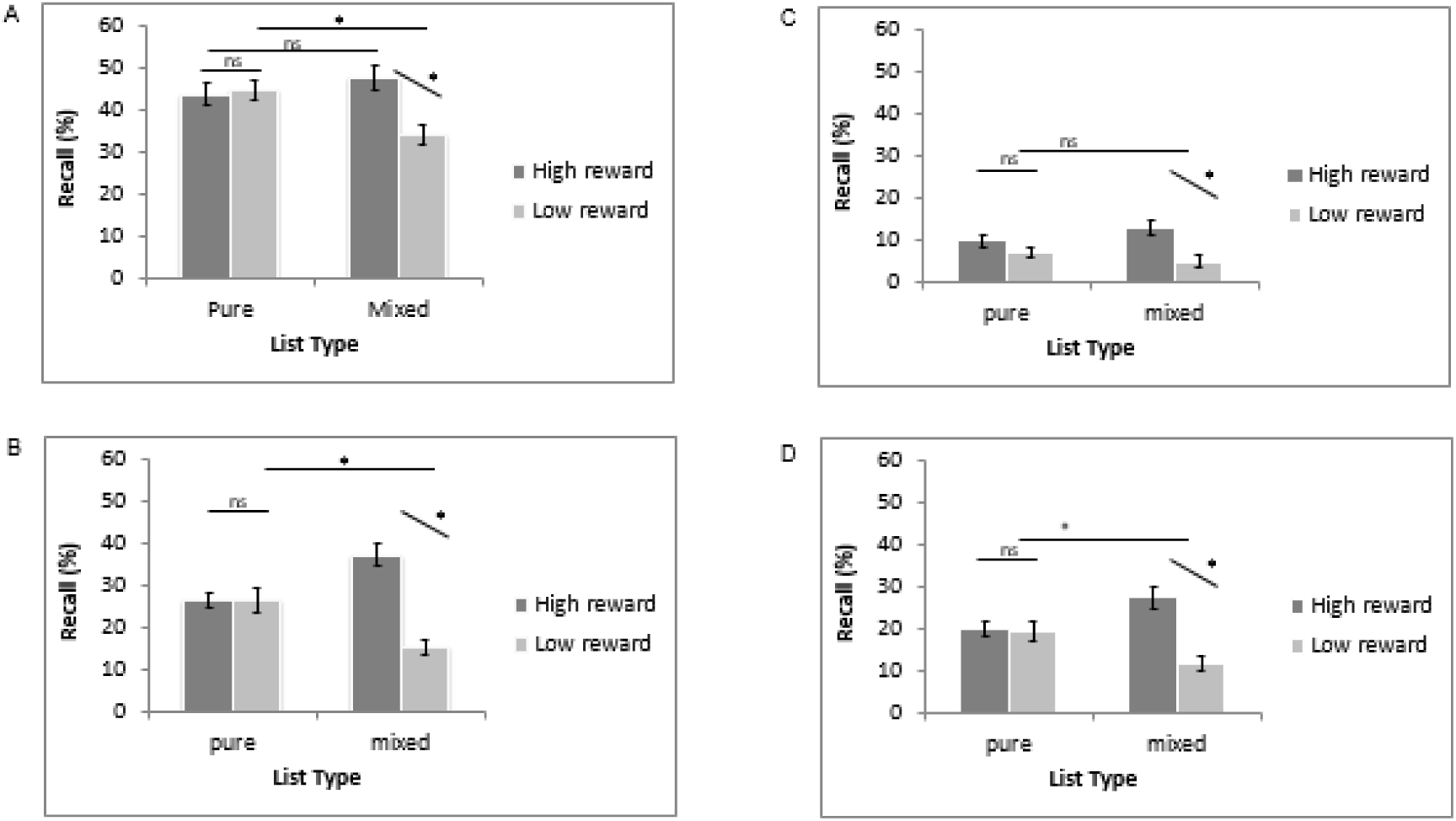
Average free recall as a function of reward and list composition. High and low reward are presented in dark and light grey, respectively. Error bars correspond to the standard error. A: Immediate memory test in Experiments 1. B: Immediate memory test in Experiment 2. C: Delayed memory test in Experiment 2 for lists that were only tested once. D: Delayed memory test in Experiment 2 for lists that were tested twice.

#### Experimental results

All Participants complied with the instruction for minimum distractor task performance. Testing memory with free recall is more straightforward when the stimuli are words, because then it is more obvious when participants did or did not recall a target. Scoring recall of picture stimuli is more open-ended.

Regardless of the specific instructions participants receive, we cannot be sure that they have retrieved more than a gist of the picture. In our previous work, we asked participants to describe a picture with a short sentence in order to distinguish it from all others. Here we required participants to provide three details about each picture in order for the response to be scored as correct. These instructions remove instances where participants only have a very weak memory of item gist. The benefit of the current instructions is that they are clearer to participants and thus decrease measurement error and inter-individual variance. In accordance with the instructions given to participants, picture recall was scored as ‘correct’ if participants recalled at least 3 correct details about a picture (disregarding participants’ own parsing of details with commas). Two raters coded the recall data independently. Although what counts as ‘detail’ was not defined absolutely, the raters shared a practical understanding of what counted as a detail, evident in high inter-rater reliability (Cronbach’s α = .96). Discrepancies in their scoring and they were resolved through discussion. Given that here, participants knew they would only be scored as correct if they retrieved 3 details, some of them may not have even attempted recall if they only had a single detail. Therefore, a score of 0 for any recall could mean either no memory at all, or memory that is not strong enough. The proportion of correct recalls was calculated for each condition (out of 16 in the two pure list conditions, and out of 8 in the mixed list condition). Items which could not be confidently assigned to the preceding list were coded as intrusions (combining unstudied items and items from other lists). Three paired t-tests compared the average intrusions for pure low-reward and pure high-reward, and each pure list to the mixed list. No significant differences were found, *t*<1.

The statistical analysis of average recall follows exactly on the analytic approach we have used in previous work. That work led to the prediction that we would find an interaction between list type and reward. Therefore, the four comparisons that ensue – within lists (high vs. low reward, in pure and mixed lists separately) and across lists (high and low reward in mixed vs. pure lists) are all planned contrasts. To be conservative we corrected these for familywise error by using a Bonferroni-corrected p-value threshold of *p*<.0125.

Average recall across experimental conditions was just under 50%, the level expected based on computational models of recall (Naim et al., 2019). The average recall of mixed lists, collapsing across reward (*M*=41.27, SD=12.48) was numerically lower than the average recall of unrewarded pure lists (*M*=43.86, SD=13.79). The average free recall data from each of the conditions are depicted in Figure 2. They were analysed with a 2 (list: pure, mixed) x 2 (reward: high, low) repeated-measures ANOVA. The effect of list type was not significant, but the interaction with reward was significant, *F*(1,28)=20.09, p<.001, η_p_^2^ =.42, qualifying the significant main effect of reward, *F*(1,28)=10.71, *p*<.001, η_p_^2^ =.28. These results reveal a reward-dependent list-composition effect. Planned paired t-tests showed that the effect of reward was only significant in the mixed list condition, *t*(28)=4.38, *p*<.001, Cohen’s d = .81. There was no significant difference between recall of pure lists, *t*<1 or between memory for high-rewarded items in mixed and pure lists, *t*(28)=1.69, *p*>.1, Cohen’s d =.38 Low-rewarded items were recalled less well in mixed lists, *t*(28)=4.72, *p*<.001, Cohen’s d =.88.

### Experiment 2

#### Simulation results

We begin by using eCMR to simulate Experiment 2. The simulations reported here use *φ_emot_* = 2.25, as in Simulation 2a, because they provided a better qualitative fit to the mixed list results of Experiment 2 than the value used in Simulation 1a. The simulation used the list structure of Experiment 2, where there is only one semantic category in each list. Simulation 3 models the immediate test results of Experiment 2. Simulation 4 decreases the ability of recalled items to retrieve the temporal context of their encoding, and Simulation 5 also decreases the ability of recalled items to retrieve the source context of their encoding. Simulation 6 did all of this but also added a diminished weighting of the temporal context in recall. All 4 operationalisations of delay resulted in a list-composition effect (Figure 3).

**Figure 3.**
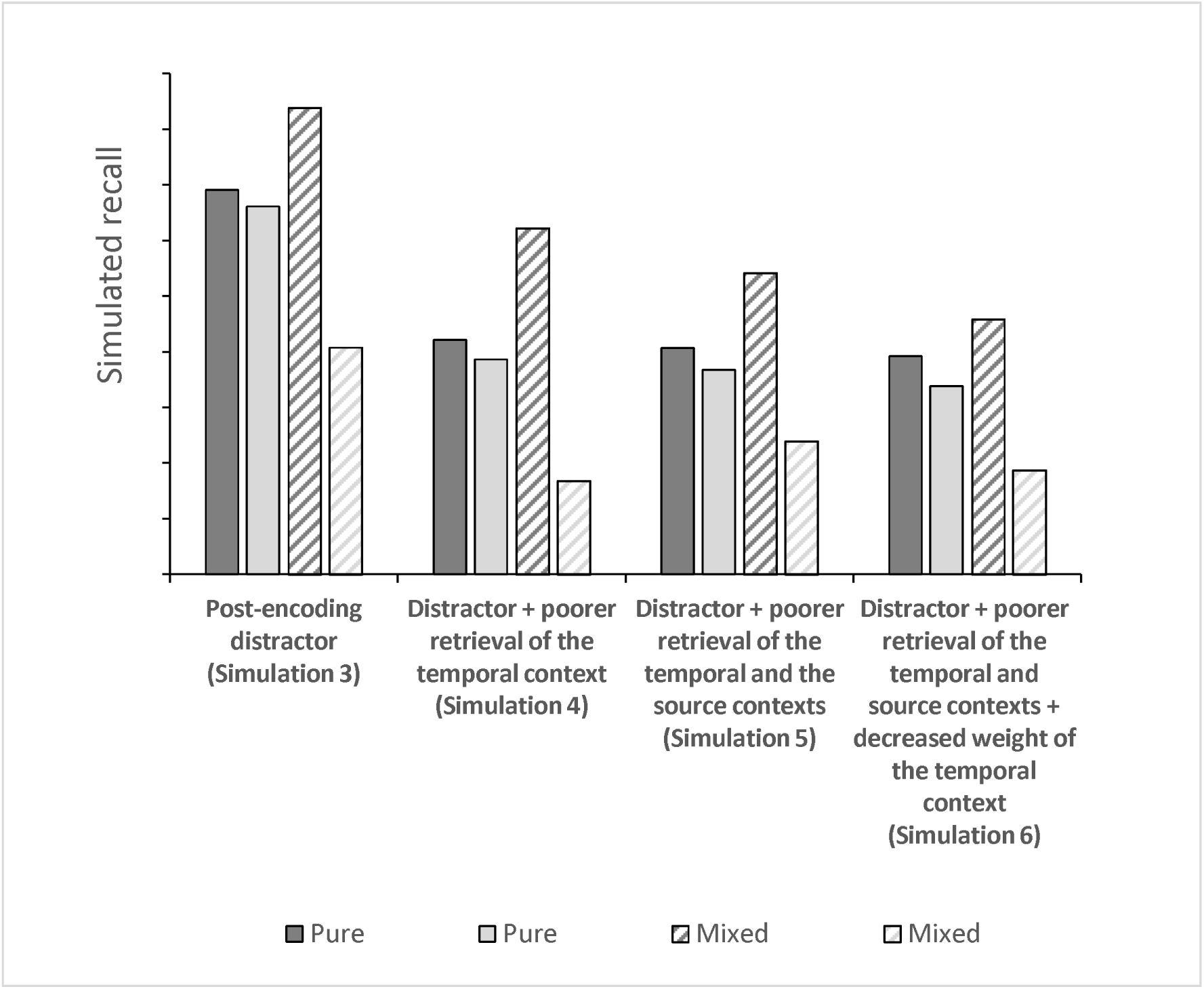
Model simulations of Experiment 2. The leftmost simulation models immediate recall. The nested simulations to the right add additional facets of delayed memory, in order: a decreased ability to retrieve the temporal context, a decreased ability to retrieve the source context, and a decreased weighting of the temporal context.

#### Distractor task performance

*Because some participants did not comply with the instructions for the distractor task, performance was analysed in more detail.* On average, participants made 0.26 errors (SD = .36) and solved 14.49 (SD = 4.62) maths problems correctly.

#### Picture recall

Recall was scored by the experimenter as ‘correct’ if participants recalled at least 3 correct perceptual or semantic details about a picture. When the experimenter (DK) identified that an experimental picture was recalled, she noted its output order and reward value in a table. Intrusions were not coded. Average immediate recall across experimental conditions was lower than that we observed in Experiment 1, potentially indicative of decreased motivation during encoding on part of this sample, or due to the faster presentation rate. Recall averaged around 26% in the immediate recall, 20% in the delayed recall of pictures already recalled in session 1, and only 9% of pictures that were only recalled for the first time in session 2. The average free recall data were analysed with a 2 (delay: immediate, delayed) x 2 (list: pure, mixed) x 2 (reward: high, low) repeated-measures ANOVA. This analysis only includes lists that were tested once – either immediately (session 1) or only after a delay (only tested in session 2), not lists that were tested in both sessions. The three-way interaction was significant, *F*(1,29)=13.83, *p*<.001, η_p_^2^=.32. Using the same analysis, but including only includes lists that were tested twice – both immediately (session 1) and after a delay (only tested in session 2), we found that as for the once-recalled data, the three-way interaction was again significant, *F*(1,29)=5.13, p<.001, η_p_^2^=.15. We unpacked these interactions to examine whether the list composition effect can be detected immediately and whether it persists after a delay.

#### Immediate recall

The first analysis was a list-by-reward repeated-measures ANOVA which focused on average recall in the immediate condition; these data are depicted in Figure 2. Its aim was to examine whether this experiment replicated Experiment 1. The effect of list type was not significant, F<1, but the interaction with reward was significant, *F*(1,29)=30.05, *p*<.001, η_p_^2^=.51, qualifying the significant main effect of reward, *F*(1,29)=20.06, *p*<.001, η_p_^2^=.41, which was also observed in Experiment 1. Planned paired t-tests showed that as in Experiment 1, the effect of reward was only significant in the mixed list condition, t(29)=6.60, *p*<.001, Cohen’s d= 1.20, where participants again recalled high-rewarded items earlier than low-rewarded items. By contrast, there was no significant difference between the recall of the two pure lists, *t*<1. To increase sensitivity to effects that may take place at the beginning of the pure list, we analysed the recall of pure lists as a function of serial position (Figure 4), but found no significant effects (reward: *F*<1; serial position: *F*(1,15)=1.43, *p*=.13; interaction: *F*<1).

**Figure 4.**
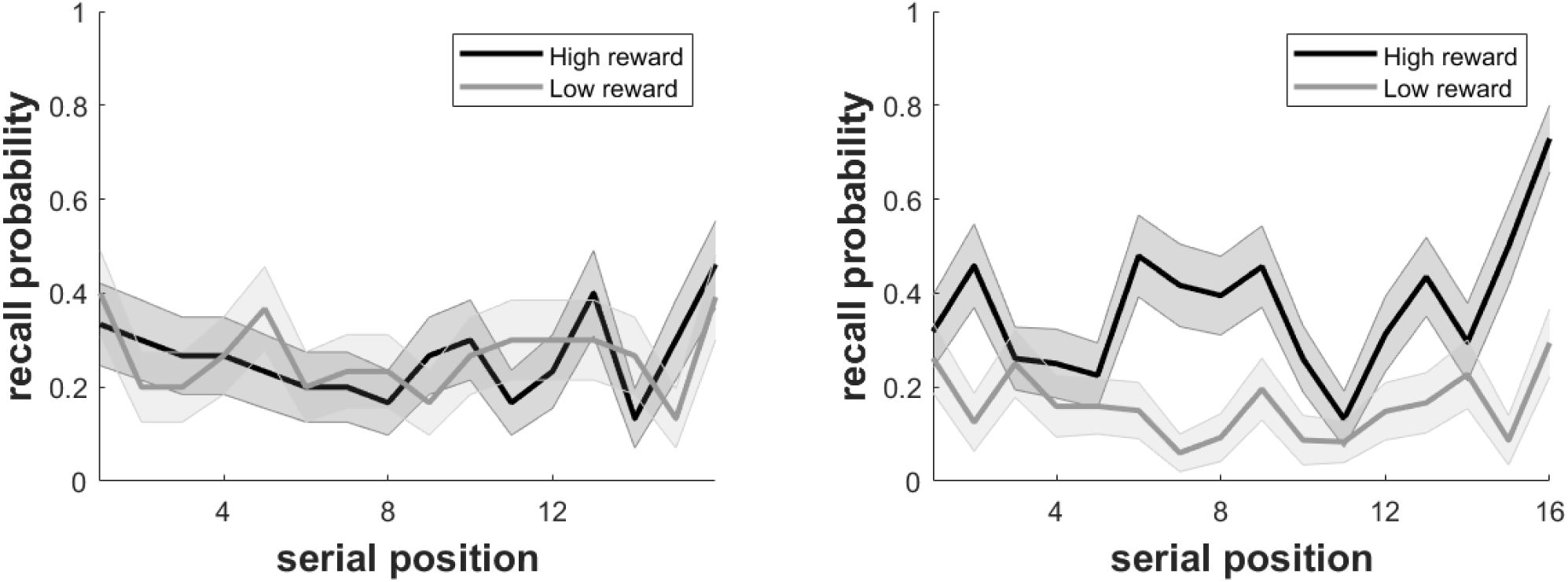
Recall probability as a function of serial position. Left. Immediate recall of pure lists in Experiment 2. The shaded area represents standard errors. Right. Immediate recall of mixed lists in Experiment 2. The shaded area represents standard errors.

As in Experiment 1, low-rewarded items were recalled less well in mixed compared to pure lists, *t*(29)=4.44, *p*<.001. However, while in Experiment 1 high reward did not increase memory, here recall of high-rewarded items was better than their recall in pure lists *t*(29) = 4.32, *p*<.001. The average recall of mixed lists, collapsing across reward, *M*=26.45, SD=10.10, was equivalent to the average recall of unrewarded pure lists, *M*=26.25, SD=9.34, t<1.

#### Delayed recall – tested once

The second analysis was a list-by-reward repeated-measures ANOVA, It focused on average recall in the delayed condition for pictures that were only recalled once (see also Figure 2). The list composition effect was less pronounced in this condition. Indeed, after a delay the interaction between list type and reward was not statistically significant, *F*(1,25)=3.40, *p*=.08, η_p_^2^ =.12, while the effect of reward was significant, *F*(1,25)=17.39, *p*<.001, η_p_^2^ =.41, and the effect of list was not significant, *F*<1. Nevertheless, planned paired t-tests showed that the effect of reward was only significant in the mixed list condition, *t*(25)=3.54, *p*<.01, Cohen’s d = .69, with a moderate-to-large effect size, but the same trend was not significant in the pure list condition, *t*(25)=1.22, *p*=0.23, Cohen’s d =.23, and had a small effect size. The average recall of mixed lists, collapsing across reward, *M*=9.61, SD=5.98, was equivalent to the average recall of unrewarded pure lists, *M*=7.93, SD=6.97, *t*(25)=1.09, *p*=.29.

#### Delayed recall – tested twice

The results of the delayed recall test in this condition mimicked the results of the immediate recall, showing that the list-composition effect persisted over 24 hours. The effect of list type was not significant, *F*<1, but the interaction with reward was significant, *F*(1,29)=22.89, *p*<.001, η_p_^2^=.44, qualifying the significant main effect of reward, *F*(1,29)=14.07, *p*<.001, η_p_^2^=.33.

#### Linear effect of forgetting

The significant three-way interaction between delay, reward and list type suggests that the number of items ‘lost’ over 24 hours differed among the 4 list types. We refer to this result as the influence of reward on linear forgetting. In order to understand this influence better, we unpack the three-way interaction with 4 separate delay-by-reward ANOVAs. The results of the analysis that included items that participants recalled once, either in session 1 or 2, and the analysis that included items that participants recalled twice, both immediately (session 1) and after a delay (only tested in session 2), yielded very similar results, and will therefore be reported together.

The analysis of pure lists found a significant effect of delay, (once-recalled: *F*(1,29)=130.07, *p*<.001, η_p_^2^=.82, twice-recalled: *F*(1,29)=48.52, *p*<.001, η_p_^2^=.63), but no other significant effects (all F<1). The analysis of mixed lists produced a different pattern. While the main effect of delay was again significant (once-recalled: *F*(1,29)=115.08, *p*<.001, η_p_^2^=.80, twice-recalled: *F*(1,29)=52.47, *p*<.001, η_p_^2^=.64), the main effect of reward was also significant, (once-recalled: *F*(1,29)=48.69, *p*<.001, η_p_^2^=.63, twice-recalled: *F*(1,29)=46.28, *p*<.001, η_p_^2^=.61), as was the interaction, (once-recalled: *F*(1,29)=13.80, *p*<.001, η_p_^2^ =.32, twice-recalled: *F*(1,29)=6.49, *p=*.016, η_p_^2^ =.18). In both cases, the significant interaction stemmed from a greater loss of high-reward items than low-reward items in mixed lists (Figure 5).

**Figure 5.**
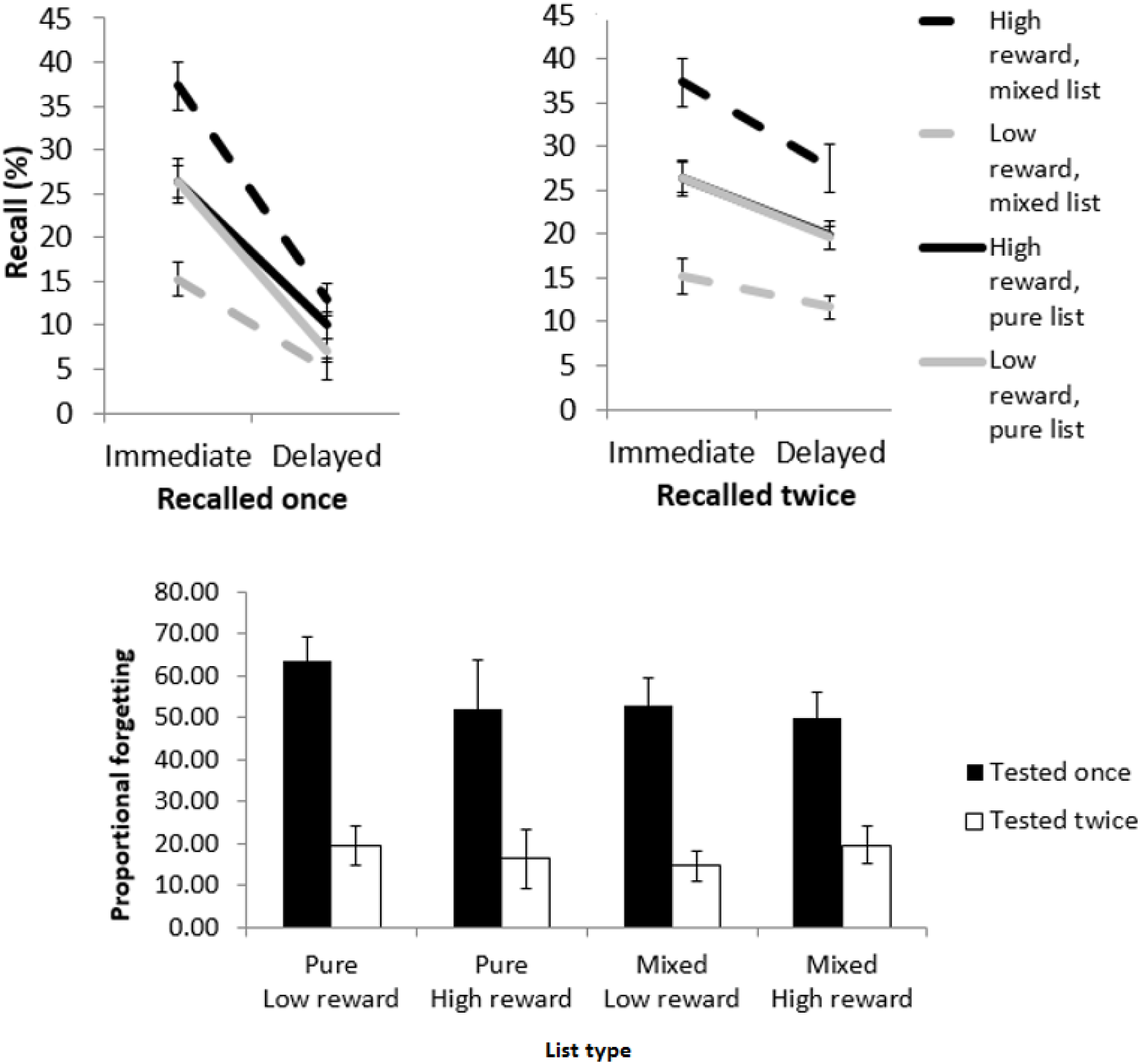
Forgetting of high and low-reward items over 24 hours. Top. Item lost over a 24-hour period. Separate plots report results for items only tested once immediately and once after 24 hours (left), or twice, both immediately and after a delay (right). Error bars represent the standard error. Bottom. Proportional forgetting scores for items only tested once immediately and once after 24 hours, or twice, both immediately and after a delay. Error bars represent the standard error.

#### Proportional forgetting rates

The number of items lost over time is a simple, intuitive metric, but it had been criticised (Loftus, 1985; Slamecka, 1985), and a variety of forgetting scores have been proposed (Wixted, 1990). Because here, immediate recall performance differed between conditions, it was imperative to examine whether reward modulated forgetting rates, taking immediate performance into account. Proportional forgetting scores were computed for each list-by-reward condition. This calculation was carried out both for lists that were tested once - immediately or after a delay, i.e. in session 1 or 2, but not both, and for lists that were tested twice, in both sessions 1 and 2. The calculation followed the formula:

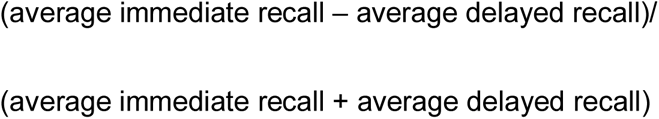

As in the d2 discrimination index in animals’ novel-object recognition tests (Ennaceur & Delacour, 1988), the denominator is a sum of both immediate and delayed recall. The benefit of this computation here is that it allowed us to avoid excluding some participants who had not recalled anything in one of these conditions. However, it still necessitated excluding participants who did not recall anything both immediately and after a delay in one of the conditions. Forgetting scores were analysed with two separate list-by-reward repeated-measures ANOVA, for the once-recalled and twice-recalled data (Figure 5). In total, *N*=26 and *N*=24 were included in these analyses, respectively. The results of both analyses were very similar: none of the factors were significant, all *F*<1.

## Discussion

In two experiments, the results of the immediate recall test showed that participants recalled significantly more high-value items than low-value items only when they were studied and recalled together, in a mixed list, but not when they were studied and recalled separately, in pure lists. This is the first demonstration of a list composition effect using an explicit reward manipulation. They replicate the emotional list composition effect, this time with emotional value operationalised as prediction of high monetary reward. The reward-based list-composition effect in average recall aligns with a core prediction of eCMR, depicted in simulations reported here and predicted by Talmi et al. (2019).

The immediate mixed-list results, where reward significantly influenced average recall, are not new. They corroborate many previous reports of enhanced immediate free recall advantage for reward-predicting items (e.g. Ariel & Castel, 2014; Stefanidi et al., 2018). Reward did not increase free recall in the immediate pure-list condition (noting that the experiments were not powered to detect small effect sizes). These results replicate previous experiments where reward, manipulated between subjects, failed to increase immediate free recall, cued recall or source memory (Murayama & Kuhbandner, 2011; Ngaosuvan & Mantila, 2005; Nilsson, 1987). Our findings suggest that those failures may be due to the between-subject nature of the manipulation of reward in those experiments, which is similar to our pure-list condition. The list-composition paradigm, which was here used with a reward manipulation for the first time, is superior to between-subject designs because it allows us to document a replicable influence of reward anticipation on immediate memory in mixed lists in the same participants. The entire pattern therefore extends the results of Ngaosuvan and Mantila and of Nilsson, where the evidence for the success of the manipulation of reward was only documented in subjective ratings of motivation. This significant interaction was replicated in both of the present experiments, and in Experiment 2 also in the twice-recalled delayed data, another advantage over a replication of null effects in between-subject designs.

An interesting aspect of our data is the comparison between the average recall in mixed lists and in what could be considered a ‘baseline’ condition, the low-reward, pure-list condition. In agreement with previous findings (Stefanidi et al., 2018), the presence of reward in mixed lists, while serving to prioritise memory for high-value items, did not increase memory for the mixed list overall, compared to baseline. This is because increased memory for high-value items was accompanied by decreased memory for low-value items in both experiments. This result has implications for the rational analysis of memory. Taken simplistically, it is always rational for an individual to recall information that has high value for them than any other information. Yet a rational agent should not slavishly recall everything that was important in the past, but rather, adapt their retrieval to present goals. Our results suggest that the goal to maximise reward may not increase retrieval capacity, but rather bias retrieval to best serve the agent’s current priorities. Retrieved-context models implement the goal to maximise reward by using the temporal context of the recall test to narrow the focus of the memory search to the most-recent list. With this narrow focus of the memory search, high-reward items stand out against low-reward items from the same mixed list, but have no particular advantage against other high-reward items. Similar ideas about the importance of the relative, rather than the absolute, distinctiveness of emotional items during memory retrieval have been reviewed authoritatively by Schmidt (Schmidt, 1991), and resemble findings from choice experiments using pain and monetary reward (Nieuwenhuis et al., 2005; Vlaev et al., 2009, 2011). These ideas could be further tested in settings where the reward landscape changes prior to retrieval.

Like Murayama and Kuhbandner (2011), we also observed that recall of pure high-reward lists compared to low-reward lists was only increased after a delay, although here the effect was not statistically significant. Reward increased the number of items lost after a delay, but had no effect on the forgetting rate. There are other reports where either emotional value or reward did not modulate the forgetting rate (Koster et al., 2015; Murayama & Kitagami, 2014), and evidence that most cognitive manipulations that increase immediate memory do not attenuate proportional forgetting rates (Loftus, 1985; Slamecka, 1985), but no conclusive meta-analysis. Murayama and Kuhbandner had double our sample size (*N*=45) and report the size of the delayed effect to be small. Our more modest sample size in Experiment 2, due to the exclusion of participants who suspected a memory test, would not have allowed us to detect a small effect, and the delayed once-recalled data of these remaining participants was low, risking floor effects. It is notable that conclusions from the once-recalled data are supported by the twice-recalled data, where floor effects are not a concern. In both analyses, the list composition effect was less pronounced after a delay, an effect that, at face value, aligns best with the hypothesis that reward attenuates forgetting. However, more high-reward items than low-reward items were forgotten in the mixed-list condition, and reward did not alter the number of items forgotten in pure lists. Neither reward, nor the type of list, modulated the proportional forgetting rate. In interpreting the null effect of reward on the forgetting rate, we note that our sample size allowed us to detect a medium effect size with >.8 statistical power. The low level of recall in the delayed test meant that many participants recalled too few items to compute various recall organisation scores, which could have helped test some of the more subtle aspects of eCMR, and possibly differentiate between the operationalisations of delay in Simulations 3-6. Tentatively, we conclude that reward does not have a large effect on forgetting in the list-composition task. Future studies can use our results to decide on a sample size suitable to detect small effect sizes. With a larger sample, it would be very interesting to compare the effect of reward on recall and on recognition memory tests, which are more prevalent in the motivated memory literature.

The replication of the list-composition effect we have obtained with emotional pictures in our previous work (summarised in Table 1 in Talmi et al., 2018) and high-reward items here is compatible with the suggestion that they impact upon the same cognitive mechanism, through prioritisation of goal-relevant stimuli for resource allocation (Mather et al., 2015; Mather & Sutherland, 2011) that then has varied influences on memory as a function of the context of recall. This is also reminiscent of findings that unsigned prediction error, possibly reflecting the initial spike of dopamine released when animals experience stimuli with either appetitive or aversive motivational salience (Schultz, 2016), drives memory enhancement (Kalbe & Schwabe, 2019). Yet although present results confirmed a core prediction of the eCMR model, we suspect that future work will reveal differences between the impact of the prospect of instrumental positive reinforcement on cognition (current manipulation) and the impact of other manipulations of emotional value, which will necessitate revisions of the model in future. For example, there are indications that recognition memory is affected differently by the influence of monetary reward compared to pain (Dunsmoor et al., 2015; Murty et al., 2016; Patil et al., 2017), and by actions taken to gain reward, compared to avoidance of action for the same purpose (Koster et al., 2015). Even within the domain of positive instrumental reinforcement, where participants act to gain reward that depends on their performance in recall tests, there is already evidence for a dissociation between actually winning and the mere possibility of winning on memory (Mather & Schoeke, 2011). It is thus important to acknowledge that even if the same model parameter is discovered to provide the best account of the impact of reward anticipation and that of negative, intense emotion, this does not imply equivalence either at the level of cognitive mechanism or at the level of neural implementation.

A limitation of the work reported here is that we did not monitor how much attention was allocated to each item during encoding. Although we hypothesized that framed items will attract more attention than unframed items regardless of the composition of the list, and this assumption is reflected in the simulations, we did not collect evidence for it. Our data allows us to test one implication of this assumption – that attention (and reward prediction error) were greater at the beginning of the high-reward pure list, compared to later in the list, conferring a special memory advantage on the early serial positions. This hypothesis was not confirmed: when we examined recall as a function of the serial position of the item in pure lists, there was no evidence for increased memory for early list items in the high-reward condition. Measuring attention at encoding could help distinguish between the theoretical account of eCMR to the list-composition effect, and alternatives such as increased rehearsal of high-reward items in mixed lists, or a form of item-method directed forgetting, where the absence of the frame serves as the forget cue and decreases rehearsal of low-reward items. While effects on massed rehearsal can be encompassed within the influence of reward on *φ_emot_* in the current eCMR framework. By contrast, eCMR currently cannot simulate selective spaced rehearsal. A memory pattern such as that reported here can stem from selective rehearsal if participants engage in spaced rehearsal of framed (but not unframed) items in the mixed list, but in the low-reward pure-list condition they engaged in spaced rehearsal of all items – perhaps because there is no cost to doing so. Such behaviour would concur with the importance of strategic effects on the value-directed remembering paradigm in Hennessee et al. (Hennessee et al., 2019), and would imply that the effect of reward is extremely sensitive to its local list context (Nieuwenhuis et al., 2005). Further research could test this alternative account either by quantifying attention during encoding, or by examining performance on a final forced-choice recognition memory test, which is more sensitive to attention at encoding, and less to retrieval biases. Increased final-recognition accuracy of high-value items regardless of list composition, together with list-composition effects in immediate recall, would align with eCMR’s predictions, but not with the selective rehearsal account. Although such a study has not been conducted with a reward manipulation, the list composition effect triggered by other manipulations is often abolished in recognition tests (Bugg & McDaniel, 2008).

In summary, our experiments show that in some situations, people recall just as much for a penny as they do for a pound. We found that memory for reward-predicting information was not any better than memory for any other information - unless the two competed during recall. eCMR predicted this pattern, and explained it as the result of the interplay between encoding and retrieval. If eCMR captures the working of participants’ minds, Sherlock Holmes may have been wrong to attribute enhanced memory for useful facts to what “the skilful workman…takes into his brain-attic” – namely, only to biased encoding. Given that retrieval capacity is more limited than storage capacity, the genius detective may have benefited also from an expert ability to select the most appropriate cues to narrow the search of his brain-attic, so that the most important facts would come readily to mind.

## Materials and Methods

### Experiment 1

#### Participants

In two previous studies, using a similar number of lists/items per list and emotional/neutral pictures (Barnacle et al., 2016, 2018), we found that the list-type-by-emotion interaction was large (the partial η^2^ was 0.33 and .61, respectively; Richardson, 2011 defines a partial η^2^>0.1379 as ‘large’). Using Gpower (Faul et al., 2007), we calculated that *N*=23 participants were required to achieve 95% statistical power to detect an interaction with a large effect size (assuming a correlation between repeated measures of 0.2). Twenty-nine participants took part in Experiment 1 (28 women, 1 man, age range 18-21). This final sample size allows us to detect a medium effect size with >0.8 statistical power. All were students, who received course credits for participation. To be eligible participants were required to have normal or corrected-to-normal vision. Participants could take part if they were between 18-35 years old, but not if they had past or current neurological or psychiatric problems, were taking centrally-acting medications, or any non-prescription recreational drugs. The project gained ethical approval from the University of Manchester Research Ethics Committee.

#### Materials

Stimuli consisted of pictures retrieved from freely-available databases, including the Bank of Standardized Stimuli (Brodeur et al., 2010, 2014) and from Food-Pics (Blechert, Meule, Busch, & Ohla, 2014; scores for pictures drawn from the Food-pic database were converted from a 1-100 scale to the BOSS 1-5 scale). 48 pictures of clothing and 48 pictures of stationery were used, for example: a green handbag, a tweed jacket, a high-heel black shoe with a buckle; a metal filing cabinet, a red paper folder. Pictures from these two semantic categories were matched for familiarity (clothing: M=4.30, SD=.32; stationery: M=4.39, SD=.35) and complexity (clothing: M=2.29, SD=.37; stationery: M=2.30, SD=.49) and adjusted in size to 280×210 pixels.

The experiment included 6 lists: 2 pure lists where all pictures were framed (pure lists, high reward), 2 pure lists where none of the pictures were framed (pure lists, low reward), and 2 mixed lists, where in each, half the pictures were framed and the other half unframed. Each list included 16 pictures, with 8 pictures from each semantic category. In the pure-list condition they were all either framed with a grey square, or unframed. In the mixed list condition, 4 pictures from each category were framed and 4 unframed, for a total of 8 framed and 8 unframed in the list. The assignment of pictures to conditions was randomised for every participant.

The experiment was implemented using Cogent200 on a MATLAB platform. In addition, the experiments used 9 printed sheets with 36 easy arithmetic subtraction and addition problems for the distractor task, with one sheet assigned randomly to each block.

#### Procedure

Participants were tested individually, in a quiet room, in the presence of a single experimenter. After providing informed consent, the experiment began with instructions and practice and ended with reward delivery. It comprised of 6 blocks, with one of the experimental lists assigned to each block. The allocation of experimental condition to block, the allocation of picture lists to the picture task in each block, the order of pictures in each list, and the order of blocks were randomised for each participant. Each block included a picture task, a distractor task, and a free recall test. These are described below.

##### Picture task

In each block participants viewed 16 pictures which could all be framed, all unframed, or half framed and half unframed. The condition of each upcoming list (pure or mixed frames) was never declared to participants. Participants did not know in advance how many framed pictures they would see in any individual block. Each picture was individually presented at the centre of the screen on a white background. It was presented for 2s followed by a blank screen for 3.5, 4 or 4.5s (randomised ISI) before the next picture was presented.

##### Distractor task

Immediately following the picture task, the screen displayed the words ‘math task’ and participants were given 60s to complete as many arithmetic questions as possible on the sheet in front of them.

##### Free recall test

At the end of the distractor task a soft beep was sounded and the words “Free Recall” were displayed. Participants were given 3 minutes to describe in writing each of the pictures they remembered at any order. They were prompted to keep trying if they stopped before the time. A beep signalled the start and end of the memory task.

##### Instructions

Before the experiment began participants were given instructions about each of the tasks. For the picture task, they were asked to view the pictures and try to commit them to memory. Participants were told that they will earn £1 for recalling framed pictures, and 10 pence for recalling unframed pictures, but they were not given any information about the proportion of framed pictures. Thus, the gain in the high-reward lists was 10 times higher than the gain in the low-reward lists. Participants were explicitly told that their goal was to maximise their monetary reward. For the Distractor task, they were informed that to qualify for reward they must (1) complete at least eight questions and (2) answer correctly two questions that are randomly selected for checking. For the free recall test, participants were asked to describe each of the pictures they could recall by writing three details about the picture, separated by a comma. They were shown two pictures and given the following examples: “A plant, spikey green leaves, in green pot” and “plastic elephant, purple, facing right”.

In previous studies of recall of complex picture scenes we did not mandate a particular number of details, but this requirement was introduced here after piloting, because of the nature of the stimuli we used, which were simpler and more inter-related than those we used in previous experiments. Indeed, some stimuli could be considered subordinates of the same basic-level exemplars (e.g. two types of shirt; Rosch, Mervis, Gray, Johnson, & Boyes-Braem, 1976), so that without this instruction, a participant might have responded with the single word “shirt” to refer to a number of list items (e.g. a folded, purple-checked shirt and a short-sleeve, blue T-shirt on a hanger). The requirement to report three details helped minimise responses that could not be confidently assigned to a particular picture. Yet we did not wish to be overly prescriptive with this instruction and therefore told participants that anything at all about the picture counted as a ‘detail’, including perceptual (the colour or orientation of an object), or semantic (an old-fashioned or an expensive-looking object). We assumed that most participants who really did recall the specific picture of the blue T-shirt would be able to describe it as a “blue (detail 1) T (detail 2) shirt (detail 3)” rather than just a “shirt”.

##### Practice

After the instructions were delivered, participants practiced each of the tasks using a set of 16 additional pictures (3 framed). During the practice block a lower reward rate was in effect (10 pence vs. 1 penny). At the end of the practice block participants were given feedback about their performance and paid any money they earned.

##### Reward delivery

At the end of the session, one block was selected randomly, and participants who qualified for reward, based on their performance on the distractor task, were paid according to their free recall performance (no participant failed to comply with the reward eligibility criteria on the distractor task). They were debriefed and thanked.

### Experiment 2

#### Comparison between experiments 1 and 2

In experiment 2 we needed to provide participants with cues that should help them, during the delayed test, to constrain their memory search to a single study list. We reasoned that participants may be confused if we ask them to recall the first, second, or third list. By contrast, it is natural to address lists by their content. Therefore, in Experiment 2 each study list included items from a single semantic category, with the category label presented before the first item of each list. Participants were tested only on half of the lists immediately. During the delayed test they were given the label of each list in turn, e.g. they were asked to recall “the list of animals”.

The added manipulation of time delay decreased the number of items participants could recall in each condition. In Experiment 1 there were 2 data points in each condition (2 pure lists in each reward condition - a total of 32 items per condition; and 2 mixed lists - a total of 16 items per condition). To avoid halving the number of items per condition and the concomitant potential increase in measurement error, we increased the number of lists from six to eight. This also allowed us to equate the number of items participants studied in each condition (now 16 in each cell). Finally, in order to decrease rehearsal effects, we shortened the ITI, introduced a cover story and a manipulation check. Other than these changes, the methodology of Experiment 2 resembled that of Experiment 1.

#### Participants

Participants in Experiment 2 were recruited from the local community through advertising. They took part in two experimental sessions and received £14 for their time and effort. We told participants explicitly that they can only qualify for reward if they do not make mistakes in the distractor task. We excluded 2 participants who made more than 2 errors on average across all lists. These participants were statistical outliers, with 5.63 and 2.25 errors, which exceeded the inter-quartile range by more than 3 SDs (the average and SDs of the performance of the final sample are provided in the results section). Of the remaining sample, 9 participants indicated they expected a memory test in session 2, and three of these 9 stated they have rehearsed the pictures in preparation for this (see “Session 2”, below). These 3 participants were excluded from the analysis to avoid contaminating the forgetting rate data. The final sample included 22 women and 8 men who were 19-35 years old. Our sample size calculations were based on the same considerations as for Experiment 1. The final sample size allows us to detect a medium effect size with >0.8 statistical power.

### Materials

Stimuli consisted of 16*9 pictures from 9 different categories: clothing, animals, cityscapes, food, household objects (non-kitchen), kitchen objects, landscapes, office items, and people. For example, a stream in an autumnal field; a pigeon on a low brick fence; a black swivel chair. They were drawn from the same sources as in Experiment 1, as well as the Nencki Affective Picture System (Marchewka et al., 2014), ImageNet (image-net.org), and internet sources. None had an overt emotional tone. While categories and pictures differed on multiple attributes, they were randomly assigned to conditions for each participant, eliminating the danger of systematic error. The Clothing list was always used in the practice block; otherwise, allocation of list to condition was randomised for each participant.

Each list included 16 pictures, which all belonged to a single semantic category. In the pure list condition, all pictures were either framed or unframed; in the mixed list condition 8 pictures were framed and 8 unframed.

### Procedure

Participants in Experiment 2 took part in two Sessions, 24 hours apart. The procedure of Session 1 was almost identical to the procedure of Experiment 1 other than the differences noted below. Session 2 included a delayed free recall test of each of the blocks encoded in Session 1.

*Session 1* included eight blocks. Each block began with a screen stating the semantic category of the pictures in that block, e.g. “landscapes”, which participants could view as long as they wished. Four blocks included pure lists (half high-reward and half low-reward), and four included mixed lists. Only half of the blocks in each condition ended with a free recall test. The ITI was eliminated, so pictures followed each other with no time gap, other than the last picture, which, due to experimenter error, followed an ITI of 3.5-4.5 seconds. As in Experiment 1, the list condition (pure, mixed, high or low reward) was never declared to participants.

#### Cover story in Session 1

Participants received the same instructions as participants in Experiment 1, and the same example pictures, but the details given with the second picture were slightly altered to: “plastic toy, elephant, purple” because the detail ”facing right” led to some participants using object orientation in their recall which were sometimes tricky to verify. Additional instructions were appended at the end. The purpose of the additional instructions was to equate the nature of encoding across blocks allocated to the immediate and delayed recall condition, and to decrease the likelihood of rehearsal between Session1 and Session 2. Participants were told that in some blocks, the immediate free recall test will be omitted. Furthermore, they were told that one of these blocks would be selected for a delayed free recall test at the end of Session 1. To maintain credibility, Session 1 always began with a bogus, additional practice block, which appeared to participants to be part of the experiment, and ended with a delayed free recall test of pictures from that block. The practice block always used the same Clothing list. In the delayed test at the end of Session 1 participants were reminded they saw a list of “clothing” and asked to recall the items from that list. Data from this test were not analysed. The session ended with reward delivery as in Experiment 1. We hoped that the inclusion of a delayed memory test in Session 1 would discourage participants from expecting another one in Session 2. Finally, participants were told that Session 2 will include different tasks in Session 2.

#### Session 2

The session began with a delayed free recall test of each of the 8 blocks presented in Session 1. Participants were shown the title of the category of a list from one block, and given 4 minutes to recall the pictures from that block, before the next title was presented. 8 titles were presented in total. Note that 4 of the tests referred to lists that were never tested before. The other 4 tests referred to lists that were already tested in Session 1, immediately after the distractor task that followed list presentation. Thus, the delayed test in Session 2 was the first test of half of the blocks, but the second test of the other half. Participants were informed that they would be rewarded for their recall according to the same schedule and rate. When participants completed the experimental tasks, they were given an additional task of picture rating. Data from this task, which are available on bioRxiv, were collected to help control stimuli in future experiments, and are not reported here. In the picture rating task, participants were asked to rate the similarity between picture pairs on a 1-7 scale. For this rating they were asked to consider picture content, rather than visual similarity, and to respond quickly. Participants were shown each category in turn (the order of categories was counterbalanced) and, within each category, all possible pairs between the 16 pictures (the order of pairs was random). In all, participants rated 120 pairs in each category, a total of 1000 ratings. At the end of the session participants were asked whether they expected a memory test, and if they answered in the affirmative, whether they made any notes or rehearsed pictures in their head. Then they were paid, debriefed and thanked.

## Acknowledgements

DT was supported by the Royal Society IE160027. DK was supported by the University of Manchester Learning Through Research initiative. ND was supported by grant DA038891 from NIDA, part of the CRCNS program, and grant 57876 from the John Templeton Foundation. We thank Charlotte Ho, Emma Kavanagh and Rebecca Lawless for collecting the data for Experiment 1.

